# SimText: A text mining framework for interactive analysis and visualization of similarities among biomedical entities

**DOI:** 10.1101/2020.07.06.190629

**Authors:** Marie Gramm, Eduardo Pérez-Palma, Sarah Schumacher-Bass, Jarrod Dalton, Costin Leu, Daniel Blank-enberg, Dennis Lal

## Abstract

Literature exploration in PubMed on a large number of biomedical entities (e.g., genes, diseases, experiments) can be time consuming and challenging comparing many entities to one other. Here, we describe SimText, a user-friendly toolset that provides customizable and systematic workflows for the analysis of similarities among a set of entities based on words from abstracts and/or other text. SimText can be used for (i) data generation: text collection from PubMed and extraction of words with different text mining approaches, and (ii) interactive analysis of data using unsupervised learning techniques and visualization in a Shiny web application.

**Availability and Implementation:** We developed SimText as an open-source R software and integrated it into Galaxy, an online data analysis platform. A command line version of the toolset is available for download from GitHub at https://github.com/mgramm1/simtext.

## 1 Introduction

Researchers utilize the primary literature to learn about the function of genes, the molecular underpinnings of disorders, and the research interests of other scientists. Frequently, groups of entities (e.g., genes, authors, disorders, etc.) are compared to another to learn about the research landscape overall. For most scientists, these efforts largely rely on time-intensive manual literature surveys. To extract higher-level information from literature in a systematic and timely manner, various tools and packages have been developed. Several R packages provide functions for data retrieval, e.g. the systematic download of abstracts from PubMed, or text mining. Without the need for programming, different web tools and databases provide summary statistics and annotations for results of a single search term (Engwall, 2017; Garcia-Pelaez *et al*., 2019; Wei *et al*., 2013) or associations and relationships among biomedical entities in the literature (Kilicoglu, 2018; Bhasuran and Natarajan, 2018; Szklarczyk *et al*., 2019; Ren *et al*., 2018).

However, such web tools and databases lack the ability to customize the analysis or visualize the results, and are focused on specific applications, e.g. on relationships among proteins. Here, we describe a semi-automatic framework for literature research: SimText. Instead of revealing associations between entities through their co-existence in single sentences and/or entire abstracts (Pavlopoulos *et al*., 2014; Szklarczyk *et al*., 2019; Junge and Jensen, 2020) we propose an alternative approach following a simple assumption: more similar or related entities share more frequently co-occurring words and scientific terms in their text than unrelated entities. SimText is a user-friendly toolset that allows users to simultaneously collect text from PubMed and extract associated vocabulary for any given set of entities. With SimText the user can interactively explore in a locally run web application key characteristics of the entities of interest and the similarities among them. This way, overlooked implicit similarities and connections in the literature can be found, and hypotheses for scientific research can be generated. All SimText tools are available as command line version (https://github.com/mgramm1/simtext) and are integrated in the online data analysis platform Galaxy (Afgan *et al*., 2018).

## 2 SimText description and workflow

SimText provides tools that can be used standalone for specific analyses or in union for a complete data analysis, from data retrieval to a web-based interactive exploration of the final results. An overview of the workflows provided by SimText is shown in **Figure 1A**. Detailed descriptions of the tools, settings, input formats, and extended versions of the use-case examples can be found in the **Supplementary Material**. SimText data analysis workflow consists of the following three modules:

**Fig. 1.**
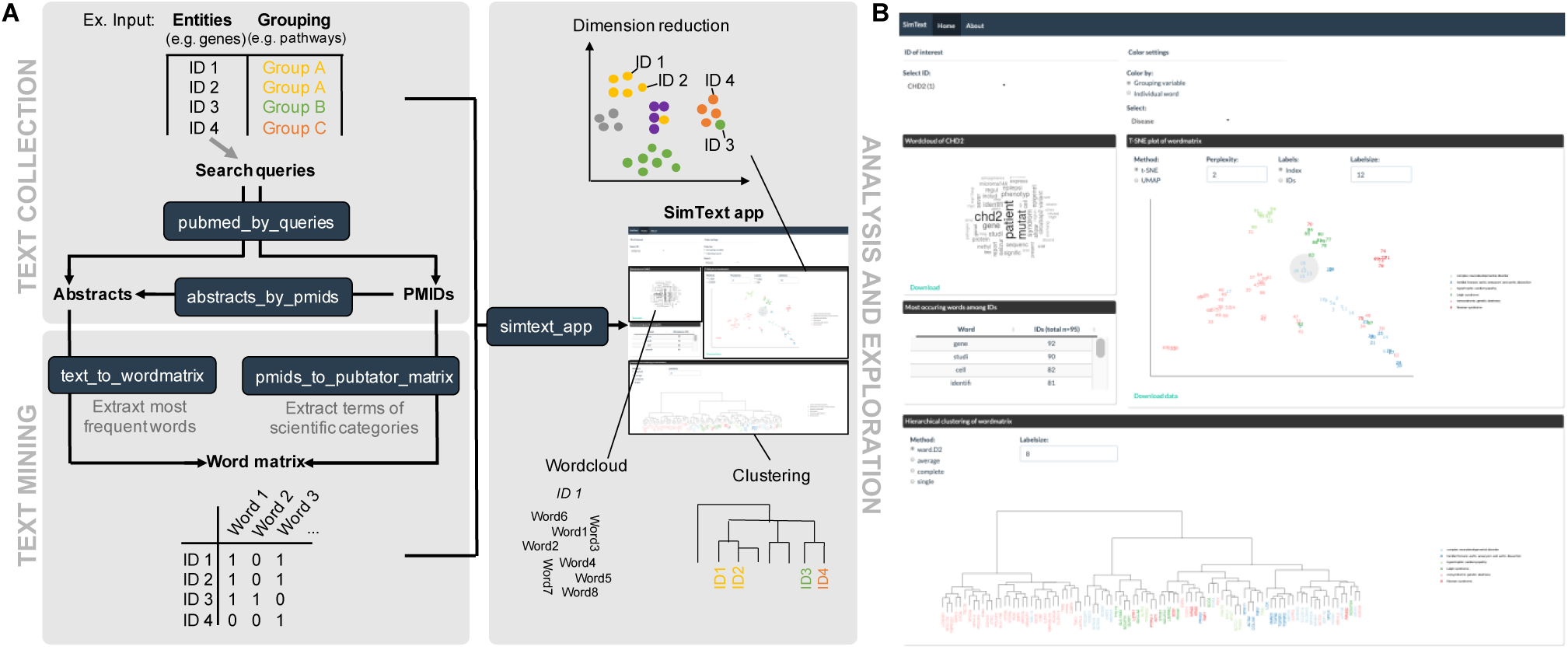
Schematic presentation of the SimText toolset. **A) Top left:** For text collection of a set of entities (e.g. gene names), the entities are used as search queries to retrieve a specified number of abstracts or PMIDs from PubMed (‘pubmed_by_queries’), or the user provides manually curated PMIDs for each entitiy that are used to fetch the corresponding abstracts (‘abstracts_by_pmids’). **Bottom left:** Given abstracts and/or additional text, associated vocabulary from each entity is extracted while providing various optional text-mining techniques (‘text_to_wordmatrix’). Alternatively, given PMIDs, scientific terms of specific categories are extracted for each entity using PubTator (‘pmids_to_pubtator_matrix’). In both approaches the output is a binary matrix with all extracted words and entities. **Right:** Interactive analysis of the generated matrix is provided by a locally run web application (‘simtext_app’). The key characteristics of the entities can be explored, and different dimension reduction and clustering techniques can be applied to the matrix to visualize similarities among the entities. Given a grouping variable (e.g. associated diseases or pathways of genes), the pre-defined grouping can be compared to the grouping of entities based on their literature. **B)** User interface of an exemplary SimText web application (https://simtext.shinyapps.io/genes).

### (1) Text collection

SimText provides two tools to collect text from abstracts related to the entities of interest (e.g. gene names). The ‘pubmed_by_queries’ tool extracts abstracts or PubMed identifiers (PMIDs) of articles of interest based on PubMed’s keyword search rules and syntax. If the user has already identified PMIDs of articles of interest, the ‘abstracts_by_pmids’ tool can be used to fetch the corresponding abstracts. In addition, the user can also provide custom text for each biomedical entity to be analyzed instead or in addition to the abstracts (for details on tools see **Supplementary Material)**.

### (2) Text mining

Using the text collected in step (1), two different types of vocabularies can be generated of each biomedical entity. The ‘text_to_wordmatrix’ tool identifies the most frequently occurring words, after word quality control, from all collected text (e.g., abstracts) for each biomedical entity (e.g., gene). Alternatively, the ‘pmids_to_pubtator_matrix’ tool extracts only scientific terms using PubTator (Wei *et al*., 2013). PubTator is a publicly available resource that performs automatic annotations of biomedical concepts (for details on tools see **Supplementary Material)**. The final output of both tools are binary matrices of all words/scientific terms and biomedical entity.

### (3) Analysis and exploration

In the last step, the generated binary matrix is used as input to a Shiny web application (‘simtext_app’). Here, among other features, the user can apply different unsupervised learning techniques (t-SNE, UMAP, hierarchical clustering) to the high-dimensional binary matrix. The user can visualize and inspect in detail groups of related biomedical entities (for details on tools see **Supplementary Material)**.

## 3 Use-case examples

As a first use-case example (for details see **Supplementary Material)**, we applied scientific knowledge about 95 monogenic disorder-associated genes and their pre-existing disorder categories from ClinGen (https://clinicalgenome.org). We hypothesized that the large-scale gene-level information extraction from abstracts could be used to visualize the similarity across monogenic disorders and possibly identify pleiotropic genes. We used SimText to retrieve 100 abstracts for each gene from PubMed, extracted the 50 most frequently used words per gene and generated a binary matrix with all words and genes (Figure S1). The app of the last step of the workflow, in which the similarity across the monogenic disorders based on literature can be explored and analyzed, can be found for illustrative purposes at https://simtext.shinyapps.io/genes or in Figure 1B.

In a second use-case example, we explored shared interests among researchers based on the word content of published abstracts. We collected names from 182 researchers from the Lerner Research Institute of the Cleveland Clinic as entities, the department they are working in as a grouping variable to identify shared interest of researchers across departments, and PMIDs of their recent publications (for details see **Supplementary Material**, Figure S1). The resulting Shiny app enables the identification of investigators with similar interest and the detailed exploration of shared terms used in abstracts (https://simtext.shinyapps.io/researchers). These use-case examples, and an additional example that uses the PubTator function, can be found in more depth in the Supplementary material. The commands, Galaxy tools and data and example files, can be found on GitHub at https://github.com/mgramm1/simtext/.

## 4 Conclusion

SimText enables the extraction of knowledge for large lists of biomedical entities (e.g., genes, authors, disorders, etc.) and the visual exploration of any existing relationships between them. Information from the literature can be automatically extracted from PubMed for each of the search queries. The processed information can be explored interactively to get insight into key characteristics and similarities among them. The SimText tools are versatile and can be used individually or in different workflows for a large number of possible use-cases. All SimText tools and can be applied to data without programming knowledge.

## References

Afgan, E. et al. (2018) The Galaxy platform for accessible, reproducible and collaborative biomedical analyses: 2018 update. Nucleic Acids Res., 46, W537–W544.

Bhasuran, B. and Natarajan, J. (2018) Automatic extraction of gene-disease associations from literature using joint ensemble learning. PLOS ONE, 13, e0200699.

Engwall, K.D. (2017) Anne O’Tate. J Med Libr Assoc, 105, 200–202.

Garcia-Pelaez, J. et al. (2019) PubTerm: a web tool for organizing, annotating and curating genes, diseases, molecules and other concepts from PubMed records. Database (Oxford), 2019.

Junge, A. and Jensen, L.J. (2020) CoCoScore: context-aware co-occurrence scoring for text mining applications using distant supervision. Bioinformatics, 36, 264–271.

Kilicoglu, H. (2018) Biomedical text mining for research rigor and integrity: tasks, challenges, directions. Brief Bioinform, 19, 1400–1414.

Pavlopoulos, G.A. et al. (2014) Biological Information Extraction and Co-occurrence Analysis. In, Kumar, V.D. and Tipney, H.J. (eds), Biomedical Literature Mining, Methods in Molecular Biology. Springer, New York, NY, pp. 77–92.

Ren, J. et al. (2018) iTextMine: integrated text-mining system for large-scale knowledge extraction from the literature. Database (Oxford), 2018.

Szklarczyk, D. et al. (2019) STRING v11: protein–protein association networks with increased coverage, supporting functional discovery in genome-wide experimental datasets. Nucleic Acids Res, 47, D607–D613.

Wei, C.-H. et al. (2013) PubTator: a web-based text mining tool for assisting biocuration. Nucleic Acids Res, 41, W518–W522.

