## Supplementary Material for "SimText: A text mining framework for interactive analysis and visualization of similarities among biomedical entities"

### 1 Description of tools

#### 1.1 pubmed_by_queries

This tool uses a set of search queries to download a defined number of abstracts or PMIDs for each search query from PubMed. PubMed's search rules and syntax apply.

Input:
Tab-delimited table with a list of search queries (biomedical entities of interest) in one column. The column header should start with "ID_" (e.g., "ID_gene" if search queries are genes).
Settings:

- Save abstracts or PMIDs
- Number of abstracts or PMIDs that should be retrieved per entity
- NCBI API key if applicable
- Name of the output file

Output:
A table with additional columns containing PMIDs or abstracts from PubMed.

#### 1.2 abstracts_by_pmids

This tool retrieves abstracts for a matrix of biomedical entities and PMIDs. The abstract text is saved in additional columns.

Input:
Tab-delimited table with rows representing biomedical entities and columns containing the corresponding PMIDs. The names of the PMID columns should start with "PMID_" (e.g., "PMID_1", "PMID_2" etc.).
Settings:

- Name of the output file

Output:
A table with additional columns containing abstract texts.

#### 1.3 text_to_wordmatrix

The tool extracts for each row the most frequent words from the text in columns starting with "ABSTRACT" or "TEXT. The extracted words from each row are united in one large binary matrix, with 0= word not frequently occurring in text of that row and 1= word frequently present in text of that row.

<u>Input:</u>
The output of ‘pubmed_by_queries’ or ‘abstracts_by_pmids’ tools, or a tab-delimited table with text in columns starting with "ABSTRACT" or "TEXT".
<u>Settings:</u>

- Number of most frequent words that should be extracted per row (default: 50)
- Convert all characters to lower case (default)
- Remove a set of English stop words (e.g., ‘the’ or ‘not’) (default)
- Transform words in the plural to their singular form (default)
- Remove any numbers in the text
- Apply Porter’s stemming algorithm: collapsing words to a common root to aid comparison of vocabulary
- Name of the output file

<u>Output:</u>
A binary matrix in that each column represents one of the extracted words.

#### 1.4 pmids_to_pubtator_matrix

The tool uses all PMIDs per row to extract "Gene", "Disease", "Mutation", "Chemical" and "Species" terms from the corresponding abstracts, using PubTator annotations. The user can choose from which categories terms should be extracted. The extracted words are united in one large binary matrix, with 0= term not present in abstracts of that row and 1= term present in abstracts of that row.

<u>Input:</u>
Output of ‘abstracts_by_pmids’ tool or tab-delimited table with columns containing PMIDs. The names of the PMID columns should start with "PMID_" (e.g. "PMID_1", "PMID_2" etc.).

Settings:

- PubTator categories that should be considered (options: Genes, Diseases, Mutations, Chemicals, Species)
- Name of the output file

Output:

Binary matrix with each column representing one of the extracted terms.

#### 1.5 simtext_app

The tool enables the exploration of data generated by ‘text_to_wordmatrix’ or ‘pmids_to_pubtator_matrix’ tools in a Shiny local instance. The following features can be generated: 1) word clouds for each initial search query, 2) dimension reduction and hierarchical clustering of binary matrices, and 3) tables with words and their frequency in the search queries.

Input:

1) Input 1:

   Tab-delimited table with

   - A column with initial search queries starting with "ID_" (e.g., "ID_gene" if initial search queries were genes).
   - Column(s) with grouping factor(s) to compare pre-existing categories of the initial search queries with the grouping based on text. The column names should start with "GROUPING_". If the column name is "GROUPING_disorder", "disorder" will be shown as a grouping variable in the app.

2) Input 2:

   The output of ‘text_to_wordmatrix’ or ‘pmids_to_pubtator_matrix’ tools, or a binary matrix.

Settings:

The user can choose different settings interactively within the app.

Output:

SimText app

#### 2 Use-case examples

We provide three examples (**Figure S1**) to illustrate the use-cases of the developed tools. The data to reproduce the example analyses, as well as the executed commands, are available online at www.github/mgramm1/simtext/. In addition to the scripts stored at Github, we implemented the toolset in the online data analysis platform Galaxy (https://usegalaxy.org).

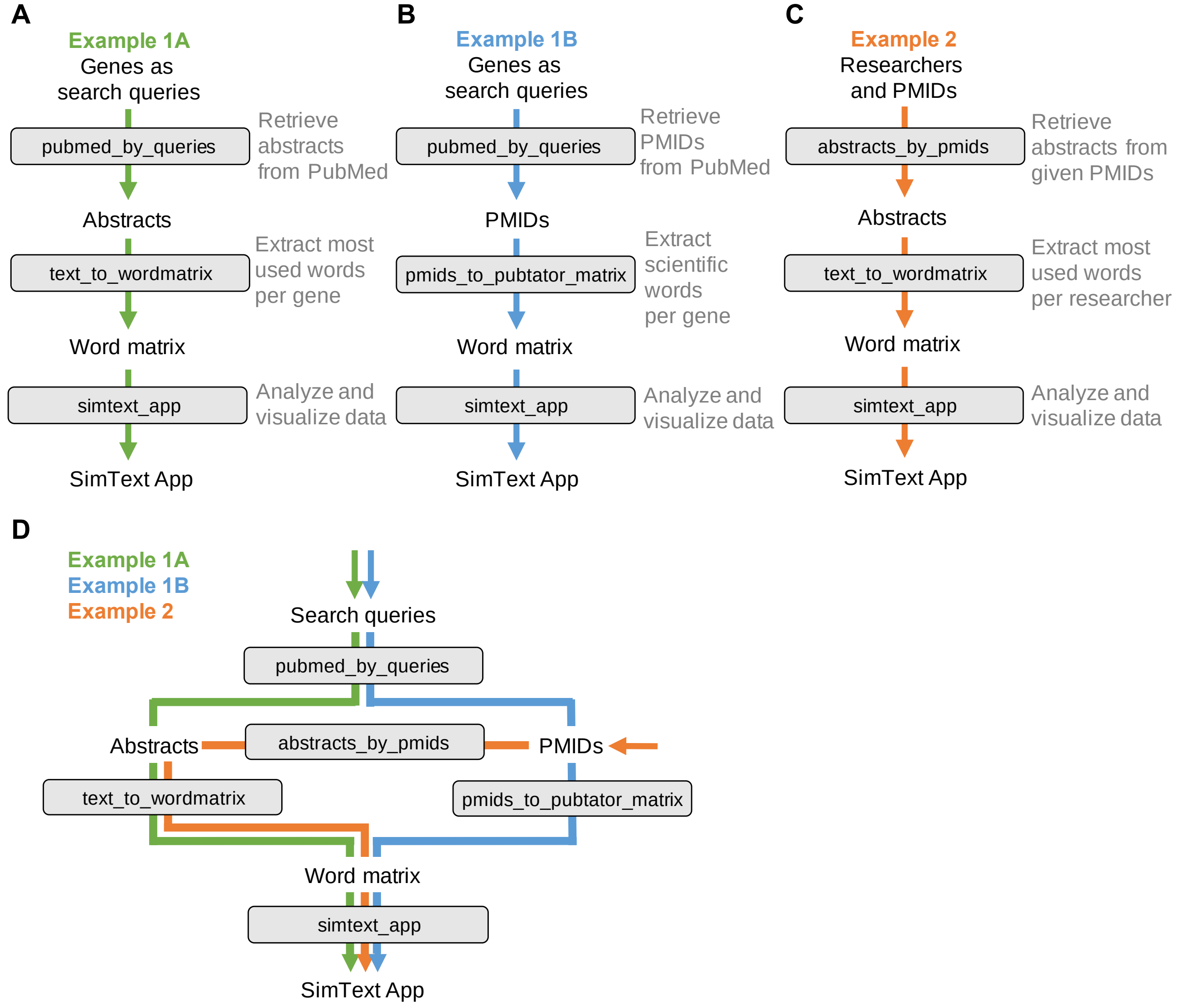

**Figure S1 Workflows of SimText use-case examples.** For detailed analysis flow of examples 1A, 1B, and 2 see below. **A)** In example 1A, we used SimText to identify pleiotropic genes by collecting information about each of the genes from PubMed and analyzing the similarities among them. We first applied the SimText tool 'pubmed_by_queries' to systematically download abstracts per gene. Next, to retrieve the key characteristics of each gene, we extracted the most frequent words from the abstracts and structured them in a word matrix ('text_to_wordmatrix'). Last, we used the SimText tool 'simtext_app' to explore the data within the interactive SimText app. **B)** In example 1B, we performed a similar analysis to example 1A. However, we extracted only biomedical concepts terms to narrow down the analysis. This was possible by the integration of the 'pmids_to_pubtator_matrix' tool in the workflow that uses PubTator, a publicly available resource that performs automatic annotations of biomedical concepts. First, we applied the SimText tool 'pubmed_by_queries' to download the PMIDs of each gene. In the next step, we applied the PubTator tool 'pmids_to_pubtator_matrix' to the PMIDs to retrieve specific scientific terms from the corresponding abstracts. Last, we used the SimText tool 'simtext_app' for interactive exploration of the data in the SimText app. **C)** In example 2, we analyzed the shared interest among researchers based on the vocabulary in their publications. First, we collected PMIDs of publications from researchers and used the SimText tool 'abstracts_by_pmids' to retrieve the corresponding abstracts. Next, similar to the methodology of example 1A, we identified the key interests of each researcher by extracting the most frequently used words from each researcher ('text_to_wordmatrix'). Last, we used the SimText tool 'simtext_app' to explore the data within the interactive SimText app. **D)** Overview of the workflows of example 1A, 1B, and 2. The SimText tools can be used in different workflows to collect data ('pubmed_by_queries' and 'abstracts_by_pmids'), extract information ('text_to_wordmatrix' and 'pmids_to_pubtator_matrix') and perform downstream analyses in the SimText app ('simtext_app').

#### 2.1 Example 1: Identification of pleiotropic genes

Rare monogenic disorders are categorized into pre-existing disorder categories. However, some gene disorders are complex and could fit multiple disorder categories. The degree of gene-level pleiotropy has not been quantified, and the lack of this information might limit the clinical understanding, and potentially, the clinical decision making to treat monogenic disorders. We hypothesized that the large-scale gene-level information extraction from abstracts could be used to visualize the similarity across monogenic disorders and identify genes that are associated with multiple disorder categories. To test this as a proof-of-concept, we selected gene-disorder groups from the ClinGen resource (https://clinicalgenome.org/). To enrich the analysis for broader disorder categories that are known for multiple monogenic disorders, we selected disorders that had more than five genes associated with it (classification strong or definitive association). This resulted in 95 genes that were associated with one out of six independent disorder categories.

##### 2.1.1 Example 1A

After curation of the gene-disorder groups, we used SimText to aggregate clinical knowledge of the genes by collecting information about each of the genes from PubMed. We applied the SimText tool 'pubmed_by_queries' to systematically download 100 abstracts per gene, totaling 9500 abstracts (**Figure S2**). Here, the tool 'pubmed_by_queries' is using the gene names recursively as search queries while PubMed’s search rules and syntax apply.

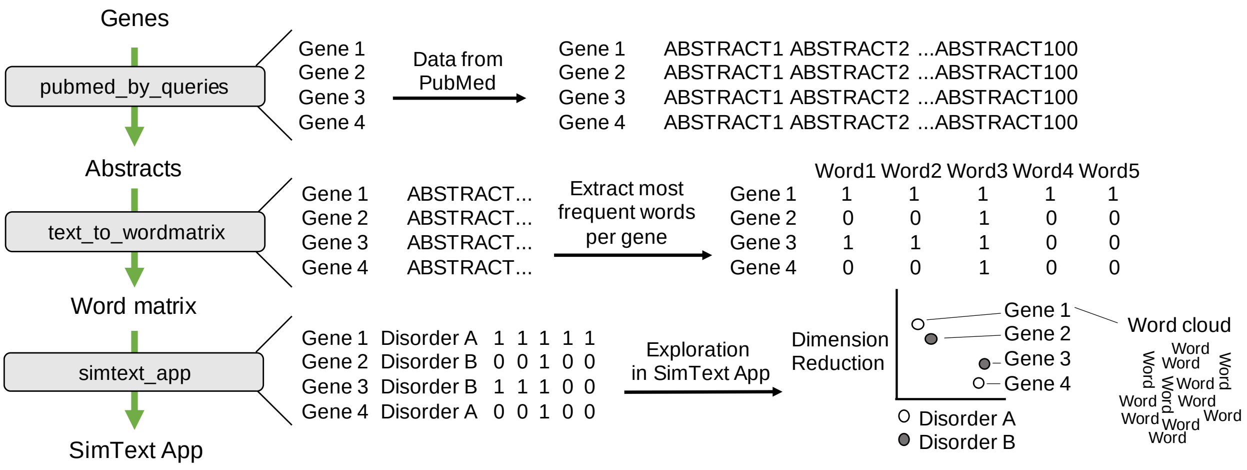

**Figure S2 Detailed workflow of example 1A.** After selecting gene-disorder groups from the ClinGen resource, we applied the SimText tool 'pubmed_by_queries' to systematically download 100 abstracts per gene. Next, to retrieve the key characteristics of each gene, we extracted the most frequent words from the abstracts and structured them in a binary word matrix ('text_to_wordmatrix'). Last, we took the gene-disorder groups (the associated disorder as a color variable) and the binary matrix from the last step to explore the data within the interactive SimText app ('simtext_app').

To retrieve the key characteristics of each gene without the need to read all the abstracts, we systematically extracted the most frequent words from the abstracts using the SimText tool 'text_to_wordmatrix'. Along with the default setting of the tool to remove English stop words (e.g. 'the' or 'not'), we selected to use the Porter’s stemming algorithm to reduce the words to its word stem (increasing the overlap of the extracted words among the genes). In this example, we set the number of words to 50, so that the 50 most frequent words per gene were extracted. The extracted words were then united in one large matrix by the tool. The word matrix, where the rows represent the genes and the columns all extracted words (see Figure S2 for details), is used to explore similarities among the genes based on counts of common words.

Next, after collecting abstracts, extracting, and structuring words for each gene (word matrix), we used the SimText tool 'simtext_app' to visualize the data within the SimText app for interactive exploration of the data. We added additional features to enable refined inspection of the data. In this example, we provided the associated disorders of the genes as a grouping variable to color code the genes in the app according to their pre-existing label. Such customization is possible for all annotations, which were provided in the original gene file (see details 'simtext_app'). In our example, this represented in addition to the gene name, the ClinGen disorder classification. For illustrative purposes, the app of example 1A can be found at https://simtext.shinyapps.io/genes/. In the example analysis and visualization, we observe that several genes do not cluster or group with genes of their pre-existing disorder categories. For example, *SMC1A* was pre-defined to be associated with complex neurodevelopmental disorders but was found in the hierarchical cluster plot (Figure S3) in a large cluster of genes all associated with genetic deafness, and in the t-SNE plot between groups of genes associated with non-syndromic genetic deafness and complex neurodevelopmental disorders. A literature search confirms the hypothesis, as mutations in *SMC1A* are associated with Cornelia De Lange Syndrome and sensorineural or conductive hearing loss (Deardorff *et al.*, 2007; Marchisio *et al.*, 2014). As a proof-of-concept, we demonstrate how the developed infrastructure could enable the generation of novel hypotheses or validate existing hypotheses in biomedical research.

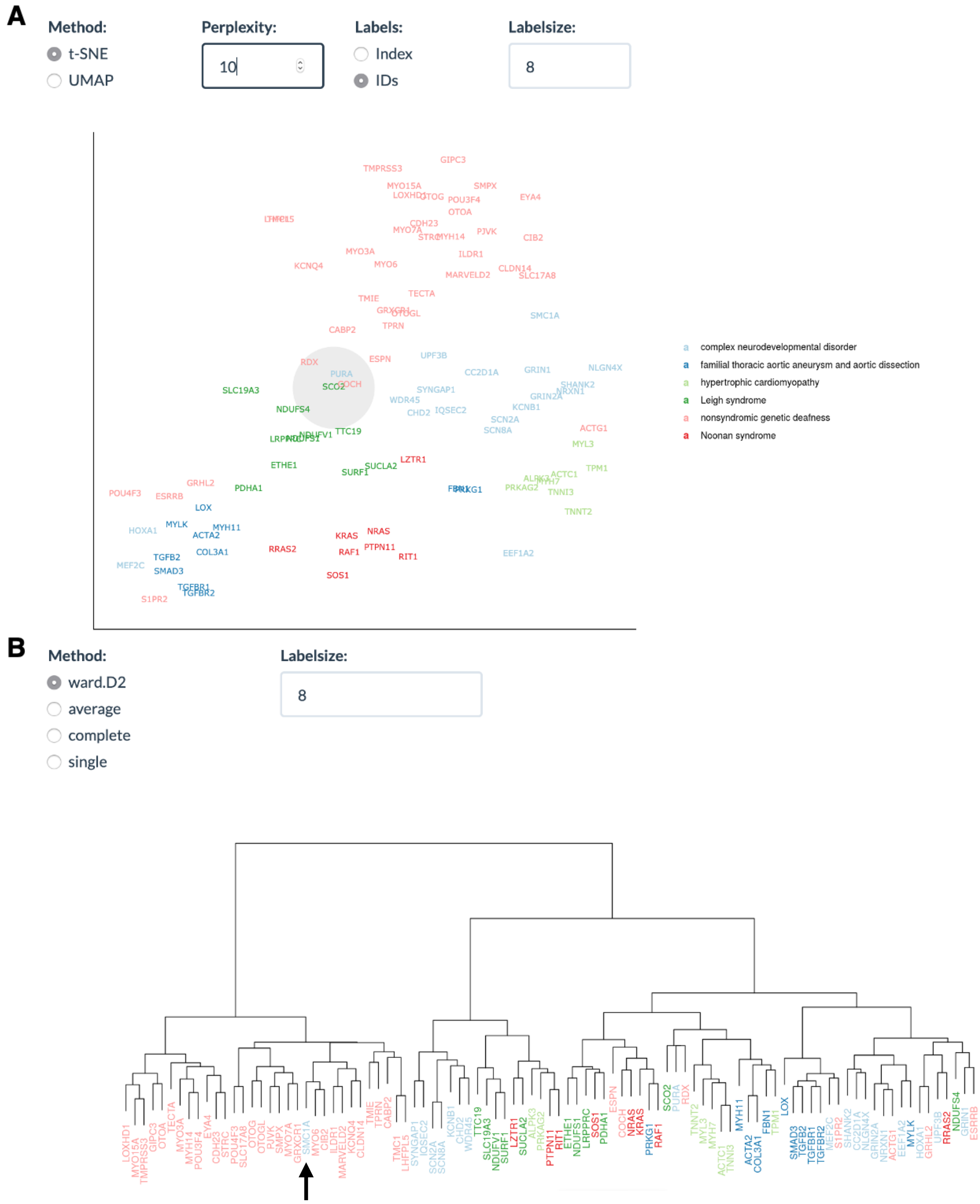

**Figure S3 Demonstration of SimText app of example 1A. A)** Two different dimension reduction techniques, t-SNE and UMAP, can be applied to the multidimensional matrix to visualize the similarities among the genes. This way, groups of similar genes can be easily identified. The color in that the genes are presented can be selected: the genes are either colored according to a grouping variable, in this case the associated disorder, or the genes are colored according to their association with one of the extracted words. **B)** In the app the user can apply different hierarchical clustering techniques to the matrix to view the hierarchy of the genes. The arrow points towards *SMC1A*, a gene we identified as a pleiotropic candidate gene (see 2.1.1).

#### 2.1.2 Example 1B

In example 1A we extracted the top 50 words across all abstracts for each gene and quantified and visualized similarities across genes. Not all frequently used words are relevant for biomedical research. Securing the quality of words can be labor-intensive. In example 1B, we performed an analysis similar to example 1A. However, we extracted only biomedical concepts terms to narrow down the analysis. This was possible by the integration of the 'pmids_to_pubtator_matrix' tool in the workflow (see **Figure S4**).

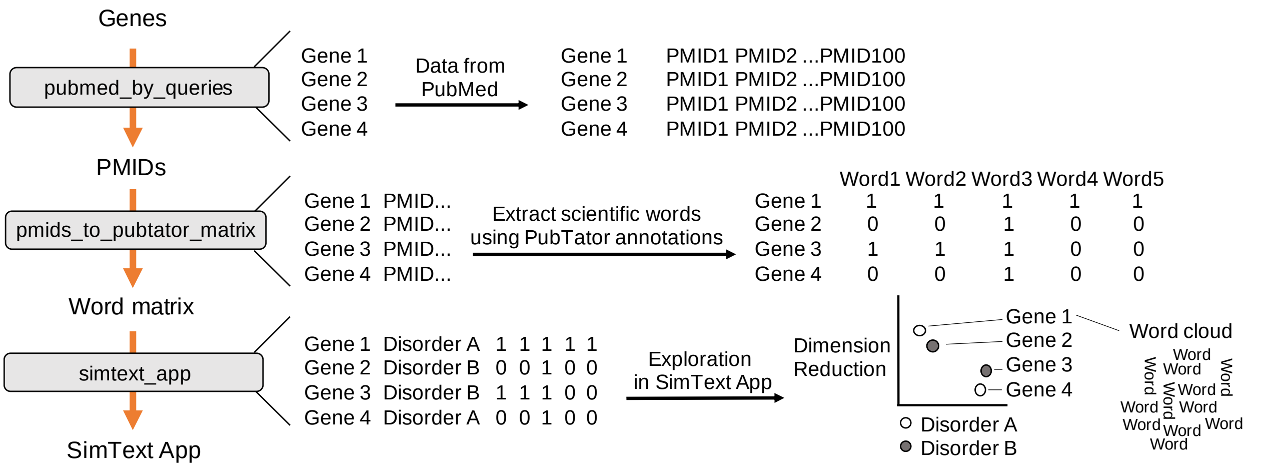

**Figure S4 Detailed workflow of example 1B.** In example 1B, we performed an analysis similar to example 1A. However, we extracted only biomedical concepts terms to narrow down the analysis. This was possible by the integration of the 'pmids_to_pubtator_matrix' tool in the workflow that uses PMIDs as input. First, we applied the SimText tool 'pubmed_by_queries' to download 100 PMIDs of each gene from PubMed. In the next step, we applied the PubTator tool 'pmids_to_pubtator_matrix' to the PMIDs to retrieve specific scientific terms from the corresponding abstracts. PubTator is a publicly available resource that performs automatic annotations of biomedical concepts. Here we selected to extract scientific terms of the PubTator categories 'Gene' and 'Disease'. By the same tool, all extracted terms from all genes are saved in one large binary matrix that enables the analysis of common scientific terms among the genes. Last, we used the SimText tool 'simtext_app' for interactive exploration of the data (gene-disorder groups and binary matrix) in the SimText app.

As a first step, we applied the SimText tool ‘pubmed_by_queries’ to systematically download 100 PMIDs per gene, totaling 9500 PMIDs (**Figure S4**). The tool 'pubmed_by_queries' is using the gene names recursively as search queries while PubMed’s search rules and syntax apply. Next, instead of manually reviewing the large number of PMIDs and their corresponding abstracts, we systematically extracted scientific characteristics of each gene with the SimText tool 'pubtator_to_matrix'. This tool takes the PMIDs from each gene and retrieves specific scientific terms from the corresponding abstracts using PubTator, a publicly available resource that performs automatic annotations of biomedical concepts (Wei *et al.*, 2013). To identify pleiotropic genes and similarities among the genes, we set the tool to extract terms of the PubTator categories 'Disease' and 'Gene'. The extracted words of each gene were then automatically united into one large matrix, in that similarities among the genes based on counts of common words can be discovered.

Analogous to example 1A, after extracting and structuring scientific words for each gene, we used the SimText tool 'simtext_app' to visualize the data within the SimText app for interactive exploration of the data.

#### 2.2 Example 2: Identification of shared interest among researchers

In large research institutions with various departments, it is difficult for researchers to know every other researcher and their research interests - even within their department. However, knowledge of shared interest can lead to fruitful collaborations and interesting research questions. We hypothesized that shared interest among 185 researchers and departments from the Lerner Research Institute of the Cleveland Clinic could be systematically assessed by their literature, or more specifically, by the identification of similar vocabulary in publications of the researchers.

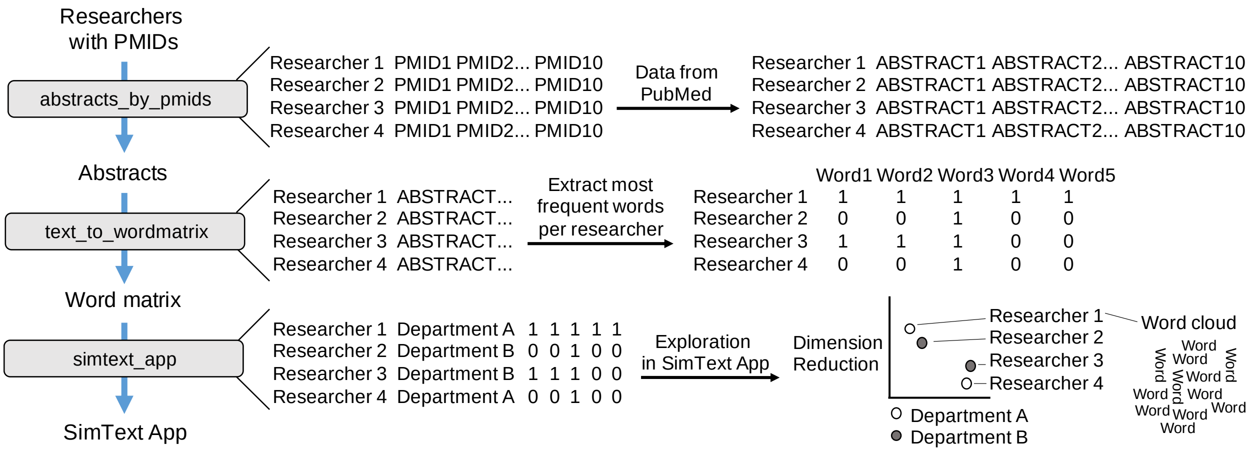

**Figure S5 Detailed workflow of example 2.** In example 2, we assessed the shared interest among researchers based on the vocabulary of their publications. First, we collected 10 PMIDs from each researcher and saved them in separate rows. We applied the SimText tool 'abstracts_by_pmids' to retrieve the corresponding abstracts. Next, similar to the methodology of example 1A, we identified the key interests of each researcher by extracting the 100 most frequently used words from each researcher (‘text_to_wordmatrix’). Last, we used the SimText tool ‘simtext_app’ to explore the data within the interactive SimText app.

In this example, we collected PMIDs of first and last author publications for each researcher as the foundation for downstream analyses similar to example 1A (see workflow **Figure S5** and in comparison to other examples in **Figure S3**). Notably, due to name ambiguity, we manually curated 10 PMIDs of publications from each researcher. To systematically retrieve the corresponding abstracts from PubMed, we applied the SimText tool 'abstracts_by_pmids' to the collected data (**Figure S5).**

Next, similar to example 1A/B, we performed the word analysis and data visualization (**Figure S3**, **Figure S5**). Following the text collection, we performed text mining to identify the shared interests of the researchers. Therefore, we extracted the most frequent words from each researcher using the SimText tool 'text_to_wordmatrix'. While applying the default settings of the tool, i.e., transforming words in the plural to their singular canonical form and removing stop words such as “and” or “not”, we selected a maximum of 100 words for each researcher. The most frequently used words of each researcher were then united in one large binary matrix.

After collecting abstracts and quantifying the words used for each researcher, we used the SimText tool 'simtext_app' to visualize the data within the SimText app for interactive exploration. Besides the generated matrix and researcher names, we provided the departments they belong to as a grouping variable to color code the researchers in the app according to their pre-existing label. The app of example 2 can be found exemplary at https://simtext.shinyapps.io/researchers/. Groups of researchers with similar interests can be identified beyond department borders by exploring the word clouds with the key interests of each researcher together with the dimension reduction plots and dendrograms (**Figure S6**).

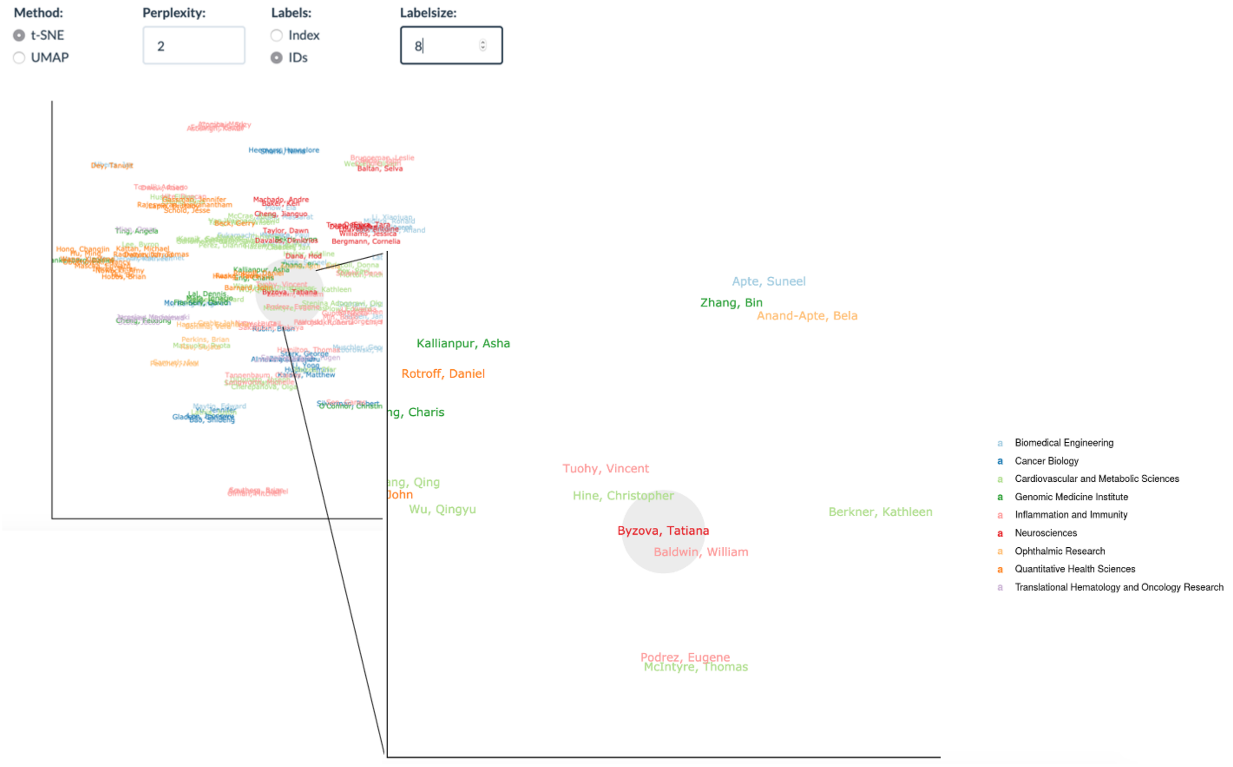

**Figure S6 Demonstration of SimText app of example 2.** The screenshot shows the t-SNE plot in that shared interest among 182 researchers can be explored. The app highlights a selected researcher of interest. The user can interactively zoom into the plot to discover potential collaboration partners.

### 3 References

Deardorff,M.A. *et al.* (2007) Mutations in cohesin complex members SMC3 and SMC1A cause a mild variant of cornelia de Lange syndrome with predominant mental retardation. *Am. J. Hum. Genet.*, **80**, 485–494.

Marchisio,P. *et al.* (2014) Audiological findings, genotype and clinical severity score in Cornelia de Lange syndrome. *Int. J. Pediatr. Otorhinolaryngol.*, **78**, 1045–1048.

Wei,C.-H. *et al.* (2013) PubTator: a web-based text mining tool for assisting biocuration. *Nucleic Acids Res*, **41**, W518–W522.
